# RAPGEF5 regulates nuclear translocation of β-catenin

**DOI:** 10.1101/152892

**Authors:** John N. Griffin, Florencia del Viso, Anna R. Duncan, Andrew Robson, Saurabh Kulkarni, Karen J. Liu, Mustafa K. Khokha

## Abstract

Canonical Wnt signaling coordinates many critical aspects of embryonic development, while dysregulated Wnt signaling contributes to common diseases, including congenital malformations and cancer. The nuclear localization of β-catenin is the defining step in pathway activation. However, despite intensive investigation, the mechanisms regulating β-catenin nuclear transport remain undefined. In a patient with congenital heart disease and heterotaxy, a disorder of left-right patterning, we previously identified the guanine nucleotide exchange factor, RAPGEF5. Here, we demonstrate that RAPGEF5 regulates left-right patterning via Wnt signaling. In particular, RAPGEF5, regulates the nuclear translocation of β-catenin independently of both β-catenin cytoplasmic stabilization and the importin β1/Ran mediated transport system. We propose a model whereby RAPGEF5 activates the nuclear GTPases, Rap1/2, to facilitate the nuclear transport of β-catenin, defining a parallel nuclear transport pathway to Ran. Our results suggest new targets for modulating Wnt signaling in disease states.

## INTRODUCTION

Canonical Wnt signaling is a highly conserved regulatory pathway for many aspects of embryonic development and adult tissue homeostasis. When dysregulated, Wnt signaling contributes to numerous diseases, including congenital malformations, neurodegeneration, diabetes, and multiple cancers (Clevers and Nusse, 2012; MacDonald et al., 2009; Moon et al., 2004; Niehrs, 2012). In the defining step of the pathway, nuclear translocation of β-catenin is essential. In fact, Wnt ligand binding inhibits the cytoplasmic β-catenin destruction machinery allowing β-catenin to accumulate and enter into the nucleus (MacDonald et al., 2009; Niehrs, 2012).

However, despite intensive investigation, the regulators of β-catenin nuclear transport have been enigmatic. β-catenin lacks a nuclear localization signal (NLS) and does not require the Ran-GTPase-regulated importin/karyopherin transport carriers to access the nuclear compartment (Fagotto, 2013; Fagotto et al., 1998; Wiechens and Fagotto, 2001; Xu and Massague, 2004; Yokoya et al., 1999). Nevertheless, there are clear parallels with the Ran based system: 1) translocation of β-catenin occurs through Nuclear Pore Complexes (NPCs), 2) the armadillo (ARM) repeats in β-catenin are structurally similar to the ARM and HEAT repeats found in importin-α and importin-β family members, and β-catenin competes for NPC-binding sites with importin-β1, 3) nuclear import is energy dependent, and 4) import of β-catenin is inhibited by nonhydrolyzable GTP analogs, suggesting the involvement of an unknown GTPase (Fagotto, 2013; Fagotto et al., 1998; Jamieson et al., 2014; Lowe et al., 2015; Xu and Massague, 2004). Together, these data demonstrate that β-catenin uses a non-classic pathway for nuclear transport; however, the molecular regulators of this process are unknown.

We previously identified the guanine nucleotide exchange factor, RAPGEF5 as a candidate disease gene in a patient with congenital heart disease and heterotaxy (Fakhro et al., 2011). Heterotaxy is a disorder of left-right (LR) patterning. Normally, our internal organs, such as the heart, are asymmetrically positioned across the LR axis, and patients with heterotaxy can have an especially severe form of congenital heart disease (Brueckner, 2007; Sutherland and Ware, 2009). However, no developmental function for RAPGEF5 had been established. Here we show that RAPGEF5, regulates left-right patterning via Wnt signaling. Importantly, RAPGEF5 is critical for the nuclear localization of β-catenin independently of cytoplasmic stabilization and the importin α/β1 mediated transport system. We propose a model whereby RAPGEF5 activates nuclear Raps to facilitate the nuclear transport of β-catenin, identifying new targets for modulating Wnt signaling in disease.

## RESULTS

### Rapgef5 is required for left – right development

To investigate whether RAPGEF5 is required for normal left-right development, we first examined organ *situs* in Rapgef5 depleted *Xenopus* embryos. The frog model, *Xenopus*, is ideal for these studies as both gain and loss of function experiments are rapid and efficient; for example, three days after manipulating fertilized *Xenopus* eggs, we can readily assess cardiac *situs* by simple inspection of *Xenopus* tadpoles (Blum et al., 2009). In vertebrates, the developing cardiac tube initially forms in the midline and then normally loops to the right (D-loop). However, defects in left-right patterning result in morphological abnormalities, including leftward (L) or ambiguous (A) loops (Brueckner, 2007; Sutherland and Ware, 2009). Knockdown of *rapgef5* using translation (R5 MO^ATG)^ or splice (R5 MO^Splice^) blocking morpholino oligonucleotides (MOs), as well as CRISPR mediated F0 depletion (Bhattacharya et al., 2015), resulted in abnormal cardiac looping and confirmed a role for Rapgef5 in left-right development (Fig. 1A). The efficacy and specificity of these knockdown strategies was confirmed by RT-PCR, western blot and RNA rescue (Fig. S1).

**Figure 1.**
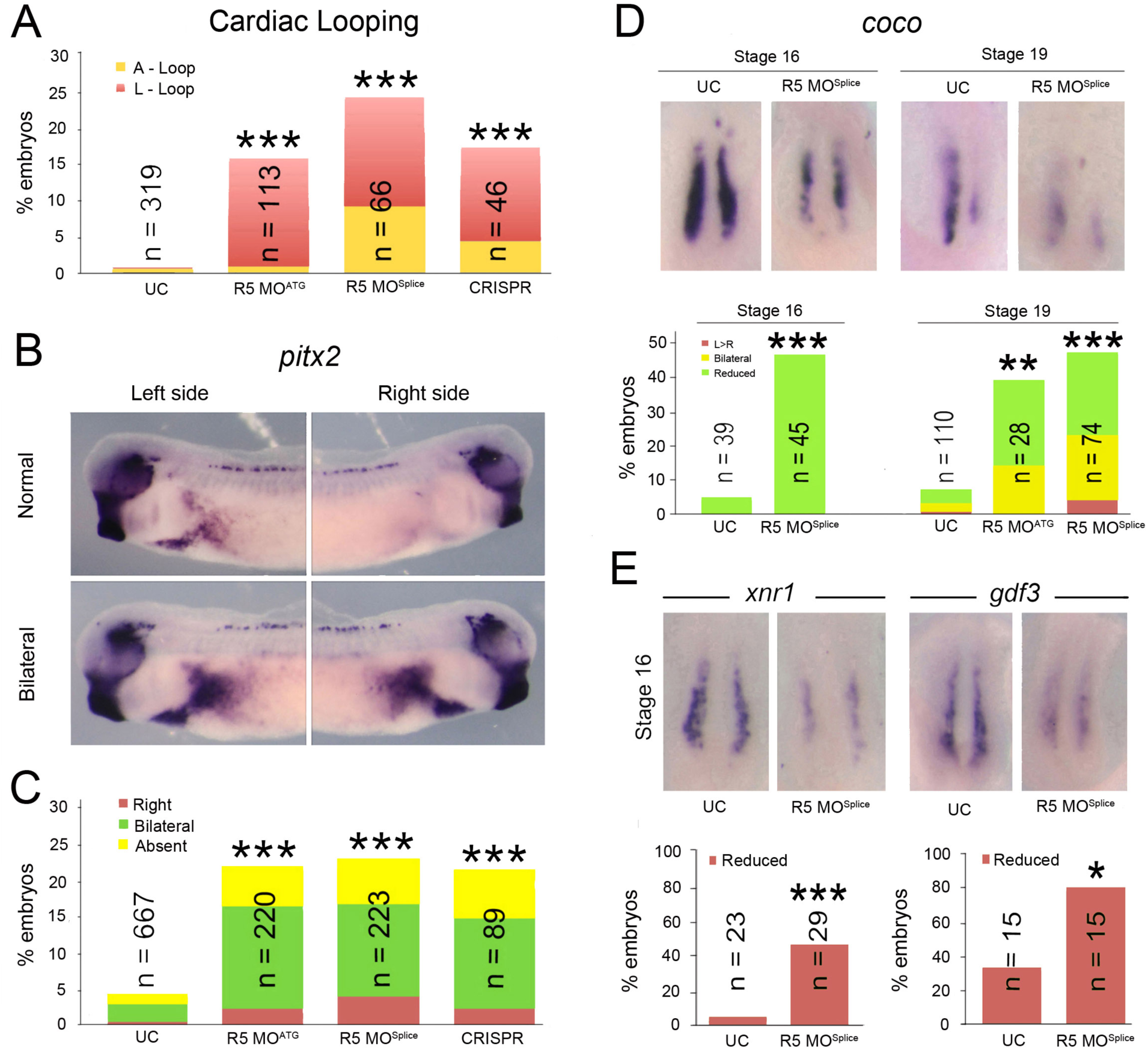
Rapgef5 depletion disrupts left-right development. (A) Percentage of Rapgef5 depleted embryos with abnormal cardiac looping (A or L loops). (B, C) *pitx2* is expressed in the left lateral mesoderm of stage 28 control *Xenopus* embryo, but is abnormally, typically bilaterally, expressed following MO or CRISPR mediated depletion of Rapgef5. (D) *coco* expression in the LRO of control and Rapgef5 depleted embryos at stages 16 and 19. Ventral view with anterior to the top. Graphs depict the percentage of embryos displaying abnormal *coco* expression. Note the reduced expression in *rapgef5* morphants at both stages. (E) *xnr1* and *gdf3* expression is reduced in the LRO of *rapgef5* morphants at stage 16, prior to the onset of cilia driven flow. Ventral views with anterior to the top. A single asterisk indicates statistical significance of P<0.05, while double and triple asterisks indicate P<0.01 and P<0.005, respectively.

Left-right symmetry is first broken at the left-right organizer (LRO). Here motile cilia generate a leftward fluid flow that is detected by laterally positioned immotile sensory cilia and translated into asymmetric gene expression (Boskovski et al., 2013; Kennedy et al., 2007; McGrath et al., 2003; Tabin and Vogan, 2003; Yoshiba and Hamada, 2014). In particular, leftward flow results in the asymmetric expression of *coco (dand5, cerl2)* in the LRO, and subsequent activation of TGF-β family members and *pitx2c* expression in the left lateral plate mesoderm (Kawasumi et al., 2011; Logan et al., 1998; Schweickert et al., 2010; Vonica and Brivanlou, 2007). Knockdown of Rapgef5 resulted in abnormalities in both *pitx2c* and *coco* expression demonstrating that *rapgef5* is required at the level of the LRO (Fig. 1B-D). Importantly in the LRO, *coco* expression is normally symmetric prior to cilia driven flow (Schweickert et al., 2010) (Fig. 1D, far left panel, Stage (st) 16); however, in morphants *coco* expression was reduced even at these pre-flow stages (st 16, Fig. 1D), suggesting a defect in the establishment of the LRO itself. This was confirmed by assaying expression of additional LRO marker genes, x*nr1* and *gdf3* (Vonica and Brivanlou, 2007), both of which were reduced before initiation of flow in morphants (Fig. 1E). Thus, Rapgef5 is required for correct establishment of the LRO and subsequent L-R development.

### Depletion of Rapgef5 impairs canonical Wnt signaling

As the LRO fails to form normally in *rapgef5* morphants, we next examined the preceding development of this structure. The Spemann organizer forms at the dorsal blastopore lip and serves to pattern dorsal structures, including the LRO (De Robertis et al., 2000; Harland and Gerhart, 1997; Khokha et al., 2005; Niehrs, 2004). There, we discovered that *foxj1* and *xnr3* transcripts, known direct targets of Wnt signaling (Caron et al., 2012; McKendry et al., 1997; Smith et al., 1995; Stubbs et al., 2008; Walentek et al., 2012), were reduced in gastrulating *rapgef5* morphants (Fig. 2A). On the other hand, numerous additional dorsal, ventral and pan-mesodermal markers were unaffected (Fig. S2). Importantly, co-injection of human RAPGEF5 mRNA rescued the *foxj1* patterning defect, demonstrating the specificity of our MO knockdown (Fig. S1D). Based on the reduction of *foxj1* and *xnr3* expression in *rapgef5* morphants, we hypothesized that Rapgef5 may play a role in Wnt signaling.

**Figure 2.**
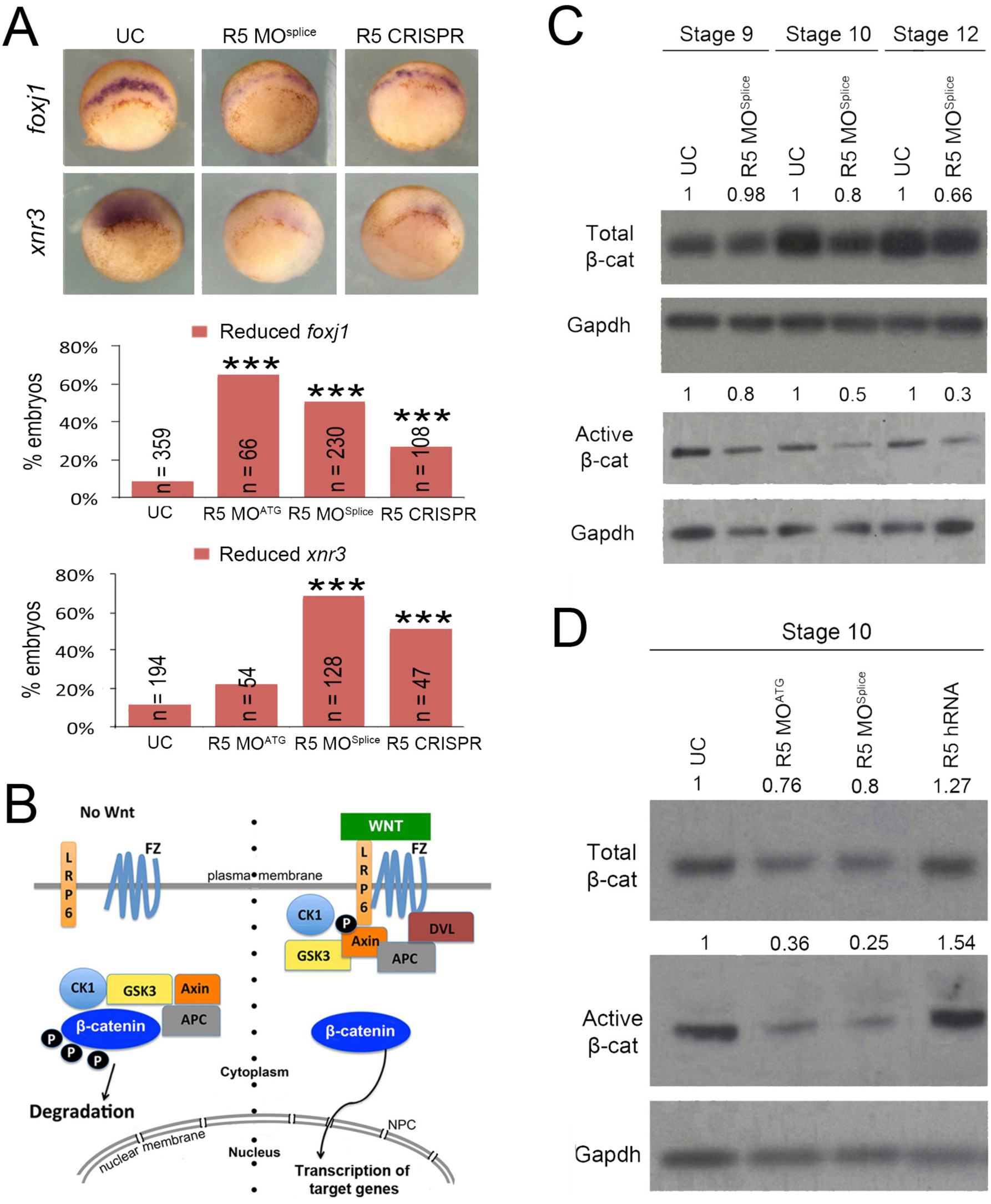
Depletion of Rapgef5 impairs canonical Wnt signaling. (A) Depletion of Rapgef5 using morpholinos or CRISPR impairs *foxj1* and *xnr3* expression in the dorsal blastopore lip of stage 10 embryos. (B) Simplified schematic of the canonical Wnt signaling pathway. Left side, in the absence of Wnt ligand, β-catenin is phosphorylated by a destruction complex containing Axin and Gsk3, which marks it for cytoplasmic destruction. Right side: Once the pathway is activated by Wnt ligand binding to Frizzled and Lrp receptors, phosphorylation of β-catenin by GSK3 is inhibited allowing β-catenin to accumulate in the cytoplasm and translocate into the nucleus to initiate transcription of Wnt target genes. (C, D) Levels of total and active β-catenin protein are essentially unchanged in Rapgef5 depleted embryos at stage 9 as assayed by western blot. However, both forms of β-catenin are reduced at stages 10 and 12. Note that levels of active β-catenin are much more severely affected. Conversely, over expression of human RAPGEF5 mRNA results in a mild increase in total β-catenin levels and a more pronounced increase in active β-catenin levels. A triple asterisks indicates P<0.005.

Wnt signaling plays diverse roles in embryonic development, including formation of primary axes, as well as in common human diseases such as cancer (Clevers and Nusse, 2012; Jamieson et al., 2014; MacDonald et al., 2009; Moon et al., 2004; Niehrs, 2012; Yamaguchi, 2001). β-catenin plays a central role in the canonical Wnt pathway (Fagotto et al., 1997; Funayama et al., 1995; Heasman et al., 1994; McCrea et al., 1993). In the absence of Wnt ligand, cytoplasmic β-catenin is phosphorylated and marked for degradation by a highly efficient destruction complex consisting of GSK3, CK1, APC and Axin (Clevers and Nusse, 2012; Davidson et al., 2005; Kofron et al., 2007; MacDonald et al., 2009; Niehrs, 2012; Taelman et al., 2010). This destruction complex maintains cytoplasmic β-catenin at low levels so that it cannot accumulate in the nucleus, repressing canonical Wnt signaling (Fig. 2B). When the Wnt ligand binds to the Frizzled receptor and the LRP6 co-receptor, the destruction complex is inactivated, and β-catenin accumulates and translocates into the nucleus to induce transcription of Wnt target genes (Bhanot et al., 1996; Davidson et al., 2005; Fagotto et al., 1997; Funayama et al., 1995; He et al., 1997; Heasman et al., 1994; MacDonald et al., 2009; Mao et al., 2001a; Mao et al., 2001b; Niehrs, 2012; Pinson et al., 2000; Tamai et al., 2000; Yang-Snyder et al., 1996) (Fig. 2B). Therefore, we began our analysis of the Wnt pathway by examining β-catenin protein levels from whole embryo lysates in *rapgef5* morphants. By western blot, β-catenin protein levels are reduced in *rapgef5* morphants at stages 10 and 12 compared to uninjected controls (Fig. 2C,D). Conversely, β-catenin levels were increased by overexpression of human RAPGEF5 (Fig. 2D). Importantly, while levels of total β-catenin were mildly affected, the active (unphosphorylated) pool of β-catenin that translocates into the nucleus was much more severely affected (Fig. 2C,D). Together these observations lead us to postulate that Rapgef5 may be required for the cytoplasmic stabilization of β-catenin.

### Rapgef5 acts downstream of cytoplasmic stabilization

To test whether Rapgef5 functions in cytoplasmic stabilization, we first asked if Rapgef5 depletion could counteract the activity of wild type (WT) and stabilized (ST) β-catenin proteins. The degradation machinery phosphorylates β-catenin at specific serine/threonine residues via GSK3; modification of these residues to alanine (S33A, S37A, T41A) prevents phosphorylation and produces a “stabilized” form of β-catenin that evades the degradation machinery (Taelman et al., 2010). Overexpression of β-catenin can induce a second embryonic axis in *Xenopus* embryos, which can be easily scored by simple inspection (Fig. 3A, left panel). Co-injection of either *rapgef5* MO or GSK3 mRNA significantly reduces the ability of WT β-catenin mRNA to induce secondary axes in *Xenopus* embryos (Fig. 3A), suggesting that Rapgef5 acts at the level of β-catenin signaling rather than upstream in the pathway. As expected, co-injection of GSK3 mRNA has no effect on the ability of stabilized β-catenin to induce a secondary axis. However, surprisingly, co-injection of *rapgef5* MO did reduce the ability of stabilized β-catenin to induce secondary axes (Fig. 3A) suggesting that Rapgef5 acts independently of β-catenin stabilization.

**Figure 3.**
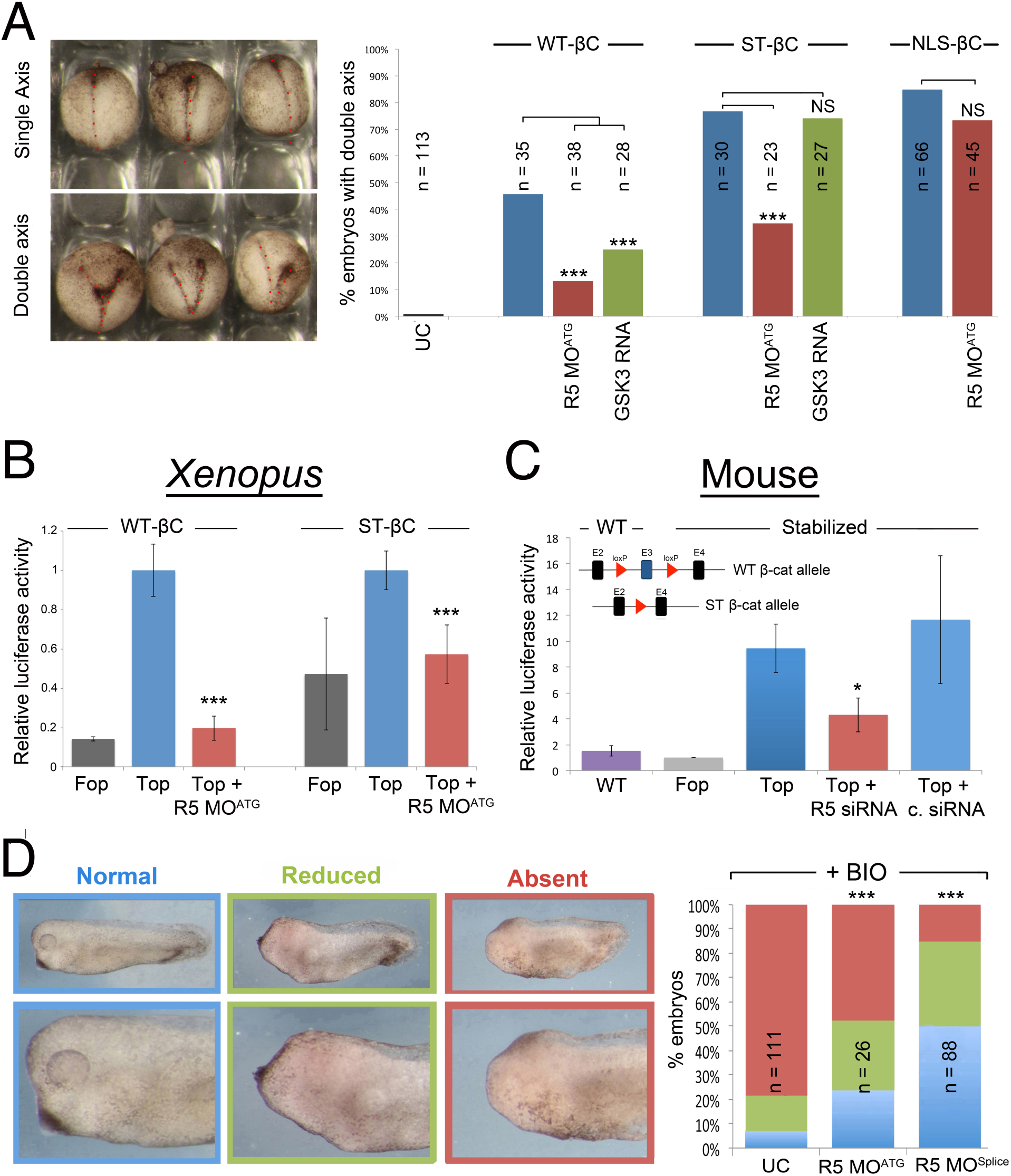
Rapgef5 acts downstream of β-catenin cytoplasmic stabilization. (A) The ability to induce secondary axes in development can be used as a read out of Wnt signaling activity. Uninjected embryos have a single axis (dotted red line) while embryos injected with β-catenin mRNA can have a second embryonic axis (two dotted lines) that are readily detected at st 16-19 embryos. Injection of WT, stabilized (ST), or NLS tagged β-catenin can induce secondary axes. Rapgef5 depletion significantly decreases the percentage of secondary axes induced by WT and ST β-catenin. Injection of GSK3 mRNA reduces secondary axes induced by WT β-catenin but has no effect on ST β-catenin. The ability of NLS-β-catenin to induce secondary axes is unaffected by reduction of Rapgef5 levels. (B) Rapgef5 knockdown reduces luciferase activity in embryos injected with WT or ST β-catenin mRNA in a TOPFlash assay. Data are represented as mean ± SD (C) Wnt signaling activity is reduced in ST β-catenin MEFs (Δexon3) following siRNA depletion of Rapgef5. The schematic depicts the *Catnblox(ex3)* mouse allele. Exon 3 (E3), containing the GSK3 phosphorylation sites that target β-catenin for cytoplasmic degradation, is flanked by loxp sites allowing for its conditional removal and production of a stabilized (ST) β-catenin allele. WT; WT cells, FOP; ST β-catenin cells transfected with FOPFlash negative control, TOP; ST β-catenin cells transfected with TOPFlash reporter plasmid, TOP + R5; ST β-catenin cells transfected with Rapgef5 siRNA and TOPFlash reporter plasmid, Top + C. siRNA; ST β-catenin cells transfected with control siRNA and TOPFlash reporter plasmid. A Renilla luciferase transfection control was included in each treatment to allow normalization. Data are represented as mean ± SEM (D) Pharmacological inhibition of GSK3 by the addition of BIO between stages 9 – 11 results in increased β-catenin signaling and loss of anterior development in *Xenopus* embryos. Depletion of Rapgef5 can counteract this effect and rescue development of the head demonstrating that Rapgef5 regulates Wnt signaling downstream of GSK3. A single asterisk indicates P<0.05, while double and triple asterisks indicate P<0.01 and P<0.005, respectively.

An alternative method to assay Wnt signaling, TOPFlash luciferase assays place luciferase expression under the control of TCF/LEF binding sites (Molenaar et al., 1996; Veeman et al., 2003). In this case, activation of Wnt signaling leads to the accumulation of nuclear β-catenin that complexes with TCF/LEF to activate luciferase expression. Consistent with our induced secondary axis experiments, Rapgef5 depletion reduced the luciferase signal of both WT and stabilized β-catenin in TOPFlash assays in *Xenopus* embryos (Fig. 3B). To see if this result was conserved to mammals, we depleted Rapgef5 using siRNAs in primary mouse embryonic fibroblasts. In mice, the specific serine/threonine residues that are phosphorylated by GSK-3 and lead to degradation of β-catenin are encoded on a single exon. Excision of this exon creates cells that express a genetically stabilized β-catenin allele (Harada et al., 1999). Even in cells with the stabilized β-catenin allele, depletion of Rapgef5 led to decreased luciferase activity (Fig. 3C).

To further test whether Rapgef5 regulates Wnt signaling independently of β-catenin degradation, we pursued a parallel strategy to inhibit GSK3 activity pharmacologically using the chemical inhibitor BIO (Sato et al., 2004). Wnt signaling must be attenuated to form the head (Glinka et al., 1998; Yamaguchi, 2001), so activation of Wnt signaling with BIO between stages 9 and 12 leads to a dramatic reduction in head development. Knockdown of *rapgef5* using either R5 MO^ATG^ or R5 MO^Splice^ could counteract the effect of BIO and rescue development of the head (Fig. 3D). Taken together in both frogs and mammals, these studies indicate that Rapgef5 regulates Wnt signaling independently of the well-established β-catenin cytoplasmic destruction machinery, suggesting that Rapgef5 acts downstream in the Wnt signaling pathway (Fig. 1B).

### Nuclear translocation of β-catenin is impaired in rapgef5 morphants

Our initial experiments with whole embryo lysates indicated that β-catenin levels were lower with Rapgef5 depletion (Fig. 2C,D); therefore, we had initially hypothesized that Rapgef5 plays a role in β-catenin degradation. Given the results described above, we sought alternative explanations and began with studies to examine β-catenin localization. To directly visualize β-catenin, we injected GFP tagged WT and stabilized β-catenin mRNA at the one cell stage and imaged the dorsal blastopore lip at stage 10. Concurrently, to enable ratiometric analysis with an importin-β1 transported cargo, we co-injected mCherry tagged with a N-terminal classic nuclear localization signal from SV40 large T antigen (“PKKKRKV”; NLS-mCherry). In control embryos, the GFP tagged β-catenin localized robustly to dorsal nuclei and the plasma membrane; however, levels of nuclear β-catenin were significantly diminished in *rapgef5* morphants whether we used the WT or stabilized variant of β-catenin. We examined this result in two ways. First, we compared GFP and RFP signals in the cytoplasm compared to the nucleus. In both WT β-catenin-GFP and stabilized β-catenin-GFP injected embryos, the nuclear:cytoplasmic ratio of β-catenin was reduced with depletion of Rapgef5 while no effect was observed with the NLS-mCherry (Fig. 4A,B far right panels). Second, we compared the nuclear signal from β-catenin (green channel) to nuclear signal of NLSmCherry (red) and observed a reduction in the β-catenin/mCherry ratio in Rapgef5 depleted embryos (Fig. 4A,B middle panels).

**Figure 4.**
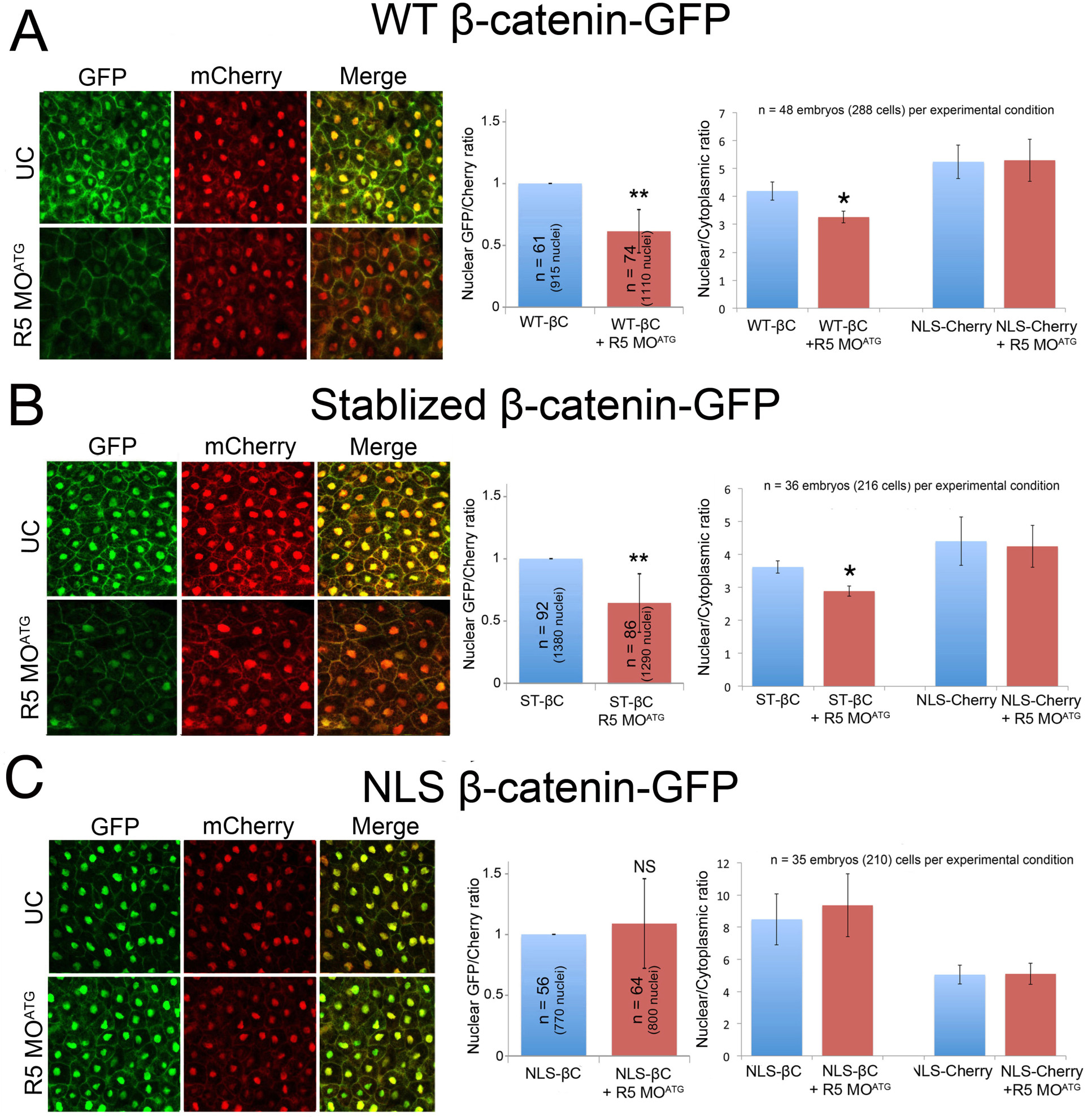
Rapgef5 is required for the nuclear localization of β-catenin. (A,B) GFP tagged WT and ST β-catenin localize to the plasma membrane and nucleus in the dorsal blastopore lip of control embryos but this nuclear localization is lost in *rapgef5* morphants (note loss of nuclear GFP signal in merged images). (C) NLS β-catenin-GFP localizes normally into nuclei even in the absence of Rapgef5. Graphs in the center panel represent the ratio of nuclear localized GFP relative to NLS-Cherry control (Data are represented as mean ± SEM). The graphs on the right display ratiometric analysis of nuclear vs cytoplasmic GFP (β-catenin) and NLS-mCherry levels. Nuclear localized β-catenin is reduced in Rapgef5 morphants relative to controls at stage 10 for WT and stabilized β-catenin but not NLS-β-catenin. Localization of NLS-mCherry is unaltered. Graphs represent mean ± SEM. A single asterisk indicates P<0.05.

We noted in these experiments that our NLS-mCherry faithfully localizes to the nucleus even when Rapgef5 is depleted and β-catenin nuclear localization is impaired. We wondered if adding the NLS signal from SV40 to β-catenin might “rescue” the nuclear localization phenotype in embryos depleted of Rapgef5. In both the secondary axis assay (Fig. 3A) and the visualization of GFP signal (Fig. 4C), addition of a classic NLS signal to β-catenin makes it insensitive to Rapgef5 depletion. Taken together, depletion of Rapgef5 has little effect on the Ran/importin-β1 nuclear transport pathway which can rescue β-catenin nuclear translocation upon the addition of a classic NLS-signal. Therefore, Rapgef5 may define a parallel nuclear transport system that is critical for β-catenin.

We next sought to evaluate β-catenin cellular compartmentalization without overexpression. We fractionated the nucleus and cytoplasm and examined the levels of β-catenin by Western blot (Fig. 5). In wild type *Xenopus* embryos at stage 10, β-catenin is present in both nuclear and the cytoplasmic/plasma membrane fractions (where it plays a critical role in cell adhesion (Brembeck et al., 2006) (Fig. 5A). In *rapgef5* morphants, levels of β-catenin were significantly reduced in the nucleus despite a relatively minor reduction in the cytoplasm (Fig. 5A,C). If we treat the embryos with the GSK-3 inhibitor, BIO, and normalize the intensities of β-catenin to control embryos without BIO treatment, inhibition of the degradation machinery increases the amount of β-catenin in both the cytoplasmic and nuclear fractions (Fig. 5B,C). However, when we deplete Rapgef5, the amount of β-catenin in the nucleus is substantially reduced despite there being more cytoplasmic β-catenin than in wildtype embryos (Fig. 5C). From these studies, we conclude that *rapgef5* depletion inhibits the nuclear localization of β-catenin even if the cytoplasmic levels of β-catenin are high due to inhibition of the cytoplasmic destruction machinery.

**Figure 5.**
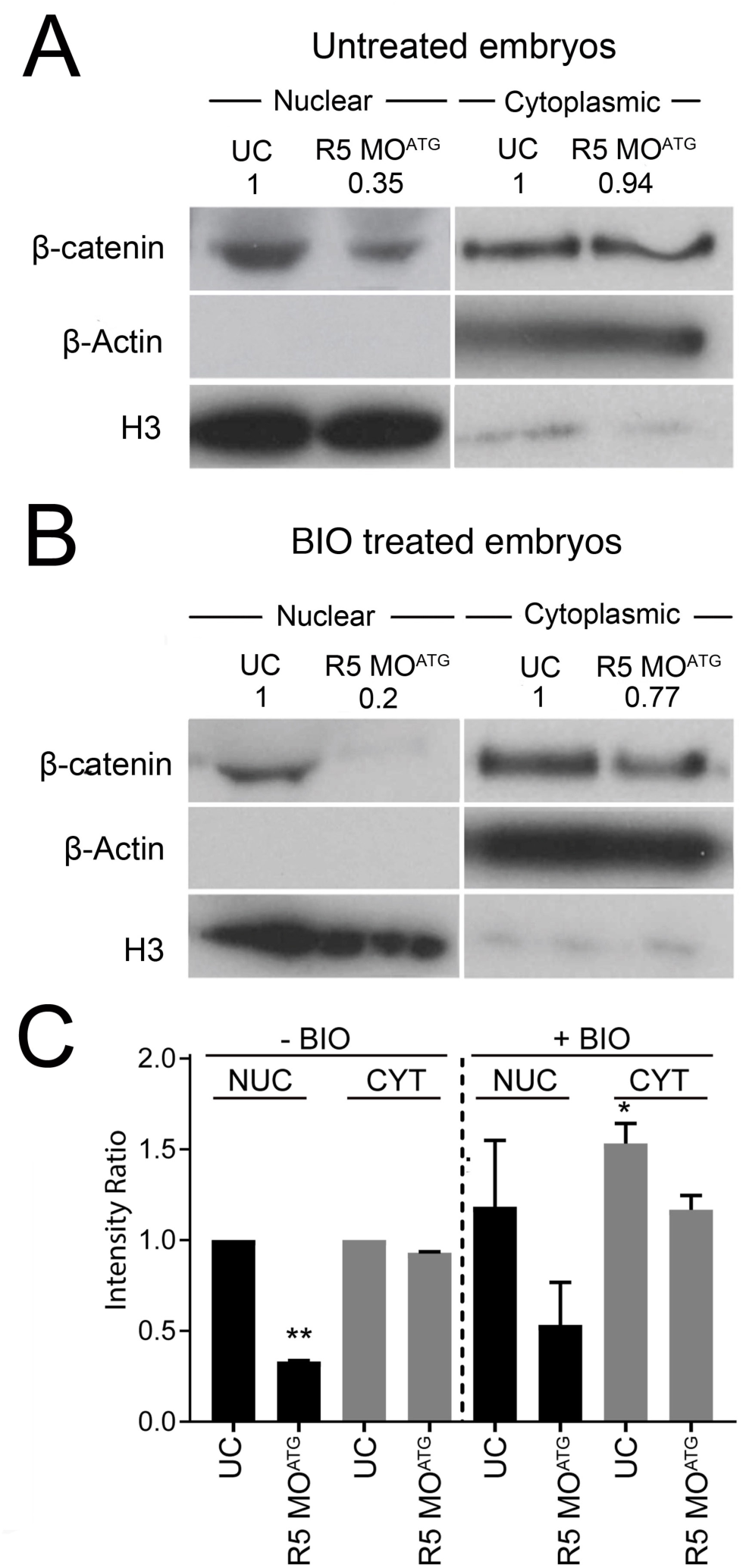
Depletion of Rapgef5 impairs nuclear translocation of endogenous β-catenin. (A) Nuclear/cytoplasmic fractionation reveals that levels of endogenous β-catenin are reduced in the nuclei of Rapgef5 morphants, while cytoplasmic levels are essentially normal. (B) BIO treatment does not rescue the nuclear localization of endogenous β-catenin. (C) Quantification of average β-catenin levels in nuclear and cytoplasmic fractions of control and morphant embryos with or without BIO treatment (error bars represent S.E.M.). All nuclear treatments (black bars) are relativized to the BIO untreated nuclear control and all cytoplasmic treatments (grey bars) are relativized to the BIO untreated cytoplasmic control. Note the reduction of nuclear localized β-catenin in Rapgef5 depleted embryos with or without BIO treatment. The BIO treated *rapgef5* morphants have less nuclear localized β-catenin than BIO untreated UC controls, despite having higher levels of cytoplasmic β-catenin. A single asterisk indicates P<0.05.

### A Rapgef5 centered nuclear translocation system

If *rapgef5* regulates nuclear translocation of β-catenin, then its expression and activity must be consistent with a role at this location. We first examined its developmental expression pattern and subcellular localization. Using anti-sense mRNA *in situ* hybridization, we find that both Rapgef5 and isoforms of its target GTPases, Rap1 and Rap2 (Quilliam et al., 2002; Rebhun et al., 2000), are expressed in multiple tissues throughout development, including the blastopore lip and LRO (Fig. S3). By immunohistochemistry, Rapgef5 protein is localized specifically to the nuclei of both whole mount and paraffin sectioned wild type *Xenopus* embryos at stages 10 and 28 (Fig. S4A) and was reduced in *rapgef5* morphants (Fig. S4A panel 2,3). Rapgef5 is also localized to the nuclei of human RPE cells and mouse embryonic fibroblasts suggesting an evolutionary conserved nuclear function (Fig. S4B,C). Consistent with previously documented roles in cell adhesion and migration (Caron, 2003), the isoforms of Rapgef5’s target Rap proteins localize near the plasma membrane as well as the cytoplasm based on immunohistochemistry or expression of fluorescently tagged proteins (Fig. S4D). Intriguingly, Rap1B, Rap2A and Rap2B also localize to the nucleus, while Rap1A accumulated in a reticular pattern that encompasses the perinuclear region suggesting a distribution throughout the nuclear envelope and ER (Fig. S4D). These results suggest an evolutionary conserved nuclear RAP system.

Based on their nuclear localization and previously reported biochemical activity (de Rooij et al., 2000; Quilliam et al., 2002; Rebhun et al., 2000), we propose a working model whereby Rapgef5 maintains nuclear Rap proteins in their active GTP bound state, which is crucial for the nuclear translocation of β-catenin. Indeed, previous reports have suggested that Rap1 localizes to the nucleus, influences nuclear β-catenin levels, and alters gene transcription profiles (Bivona and Philips, 2005; Lafuente et al., 2007). As a preliminary test of this model, we asked if nuclear Raps are in their active GTP bound conformation. GFP-^RBD^RalGDS is an active Rap sensor, consisting of the Ras binding domain of RalGDS fused to GFP, and binds specifically to the active form of Rap1-GTP and Rap2-GTP but not the inactive GDP form to allow visualization of subcellular localization (Bivona and Philips, 2005; Liu et al., 2010). In addition to previously described signals at the plasma membrane, we confirmed high levels of active Raps within *Xenopus* nuclei (Fig. 6A). Our model would also suggest that active Raps and β-catenin physically interact. In support of this, we found that Rap1 coimmunoprecipitated with β-catenin, and importantly that β-catenin is pulled down specifically with ^RBD^RalGDS, indicating that active Rap1/2 binds β-catenin in stage 10 *Xenopus* embryos (Fig. 6B,C), results consistent with a previous report in human cancer cell lines (Goto et al., 2010). Taken together, we conclude 1) Rapgef5 and multiple Rap proteins localize to the nucleus, 2) at least a subset of the Raps in the nucleus are in the active GTP form, and 3) active Rap proteins physically interact with β-catenin.

**Figure 6.**
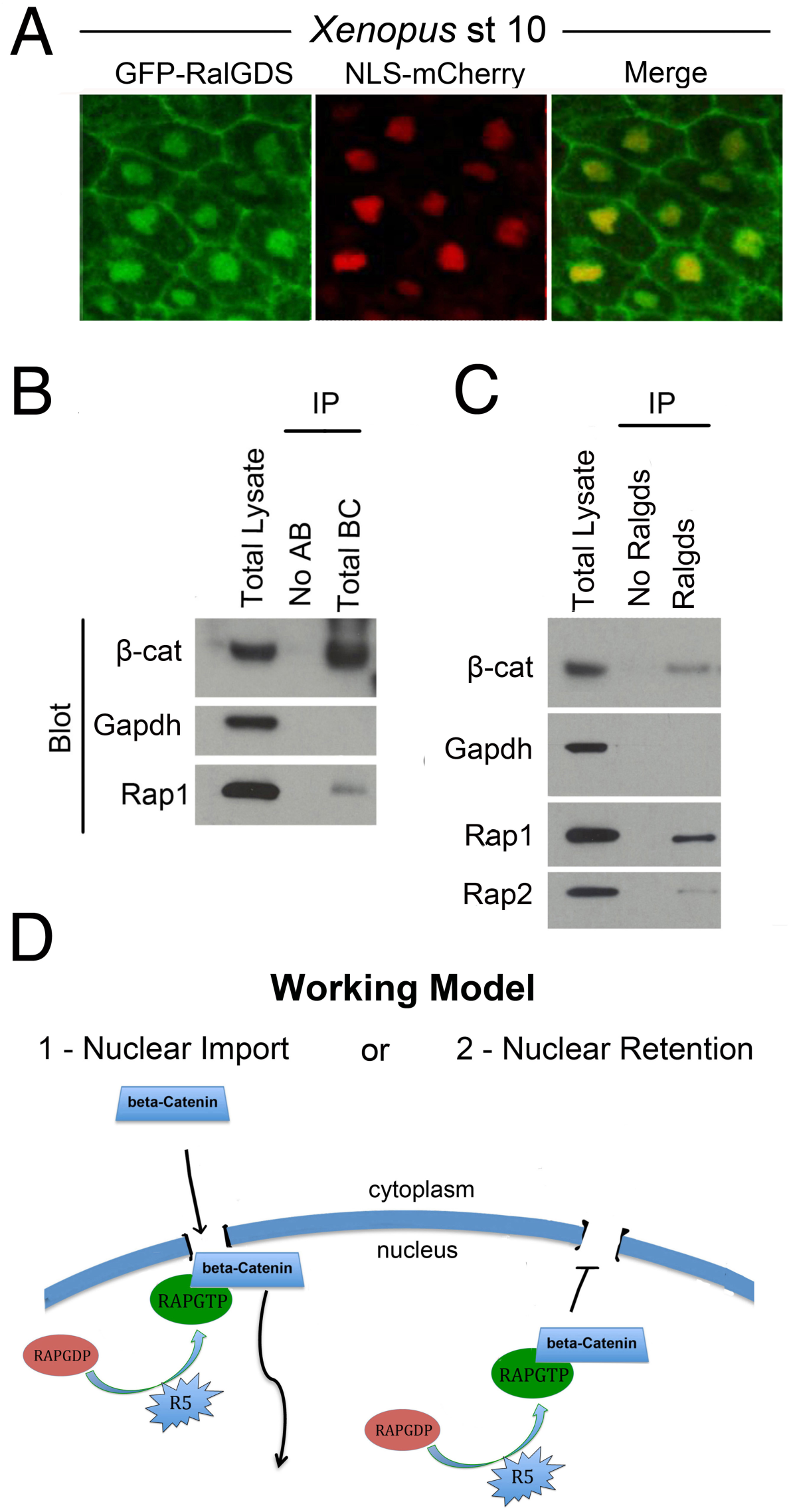
Active Raps and β-catenin interact. (A) The GFP-^RBD^RalGDS active Rap sensor reveals the presence of active Raps in the nucleus and plasma membrane of dorsal blastopore lip cells at stage 10. (B) Rap1 protein co-immunoprecipitates with β-catenin. (C) Specific immunoprecipitation of active Rap proteins from stage 10 *Xenopus* embryos using the RalGDS sensor construct. β-catenin immunoprecipitates with active Raps demonstrating an interaction, BC; β-catenin (D) Proposed model for Rapgef5s role in nuclear localization of β-catenin. Rapgef5 maintains nuclear Raps in their active GTP bound confirmation, which is required for 1) – the nuclear import of β-catenin or 2) – retention of β-catenin within the nuclear compartment.

## DISCUSSION

Integrating the sum of our data, we develop a model where RAPGEF5 activates nuclear Raps, which are necessary for the translocation of β-catenin to the nucleus. This model parallels the classic Ran based transport system, which maintains a nuclear pool of Ran-GTP through the activity of a chromatin-bound Ran-GEF, RCC1 (Kutay et al., 1997; Lange et al., 2007; Lowe et al., 2015; Xu and Massague, 2004). Structurally, β-catenin’s 12 Arm repeats are similar to the HEAT repeats found on importin-β family members suggesting that it might directly interact with the FG-nups of the NPC central channel (Fagotto, 2013; Xu and Massague, 2004). Consistent with this idea, importin-β1 can inhibit the translocation of β-catenin into the nucleus suggesting competition for NPC binding sites (Fagotto, 2013; Fagotto et al., 1998; Kutay et al., 1997; Xu and Massague, 2004). Finally, like the Ran based transport system, β-catenin’s nuclear translocation is energy dependent and depletion of ATP/GTP or addition of nonhydrolyzable GTP analogs (Fagotto et al., 1998) inhibits nuclear import, suggesting the involvement of a GTPase.

However, β-catenin nuclear import is independent of the Ran based transport system, and the identity of the GTPase is unknown. We propose that the requirement of RAPGEF5 for β-catenin nuclear translocation implicates a Rap GTPase in this role (Fig. 6D). Supporting this idea is our observation that addition of a classic NLS signal to β-catenin decouples β-catenin import from Rapgef5 and allows it to use the importin/Ran system for nuclear translocation. Also, active Raps are able to bind β-catenin and are located in the nucleus. Elucidating the precise mechanism requires further study and additional differences with the Ran based transport system may be discovered. Currently, whether Rapgef5 activity contributes to the translocation of β-catenin through NPCs or functions in a pathway that mediates β-catenin retention in the nucleus remains uncertain (Fig. 6D). Additionally, Ran-GTP acts to release karyopherin/importins from their NLS-bearing cargos; whether a β-catenin cargo is present or not remains to be seen. Importantly, Rapgef5 is localized to the nuclei of all examined cells in the embryo suggesting that its role in cell signaling is equally broad. Further studies will need to address whether the function of Rapgef5 is restricted to β-catenin or if it also effects the nuclear localization of other ARM repeat bearing proteins that lack an NLS, such as APC, p120 catenin, and Smad2.

We began our studies investigating a disease candidate gene identified in a patient with Heterotaxy and congenital heart disease. By using our high-throughput animal model, *Xenopus*, we identified an LR patterning defect in Rapgef5 depleted embryos, and given our desire to understand the pathophysiology of our patient, we discovered a novel role of Rapgef5 in the nuclear localization of β-catenin, an essential step in the Wnt signaling pathway. Wnt signaling has a broad impact across embryonic development, stem cells, and multiple diseases. In particular, aberrant Wnt signaling and Rap activity are associated with numerous cancers. Of note, increased levels of nuclear Rapgef5 have recently been linked with hereditary nonpolyposis colorectal cancer, a tumor with a well-established connection to the Wnt pathway (Chen et al., 2013). Thus, further exploration of Rapgef5 and its effectors may provide novel therapeutic targets and be of considerable relevance to human disease.

## AUTHOR CONTRIBUTIONS

JNG, FdV, ARD, and MKK conceived and designed the embryological experiments in *Xenopus.* JNG and KJL designed experiments in mouse embryonic fibroblasts. AR screened RAPGEF5 for biological activity in *Xenopus*. JNG, FdV, ARD, AR and SK performed the *Xenopus* experiments. JNG performed mouse experiments. JNG and MKK wrote the manuscript, which was read and approved by all authors.

## ACKNOWLEDGMENTS

We are grateful to the patients and families who inspired this work. Thanks to S. Kubek and M. Slocum for animal husbandry and to G. Griffin, W. Barrell, S. Gonzales Malagon, S. Rao and members of the Khokha, Liu and Gallagher labs for technical assistance and advice. We thank P. Lusk for critical comments on the manuscript. Thanks to the Center for Cellular and Molecular Imaging at Yale for confocal imaging. We also thank Prof. Philip Stork at OHSU, Prof. Han at the Pohang University of Science and Technology, South Korea and Prof. Mark Philips at NYU for providing plasmids and to Xenbase and Addgene for providing access to many of the bioinformatic resources and reagents used in this work. This work was supported by the NIH (R01HD081379 to MKK) and BBSRC (grant BB/1021922/1, BB/E013872/1 to KJL). MKK is a Mallinckrodt Scholar.

## FIGURE LEGENDS

**Figure S1.**
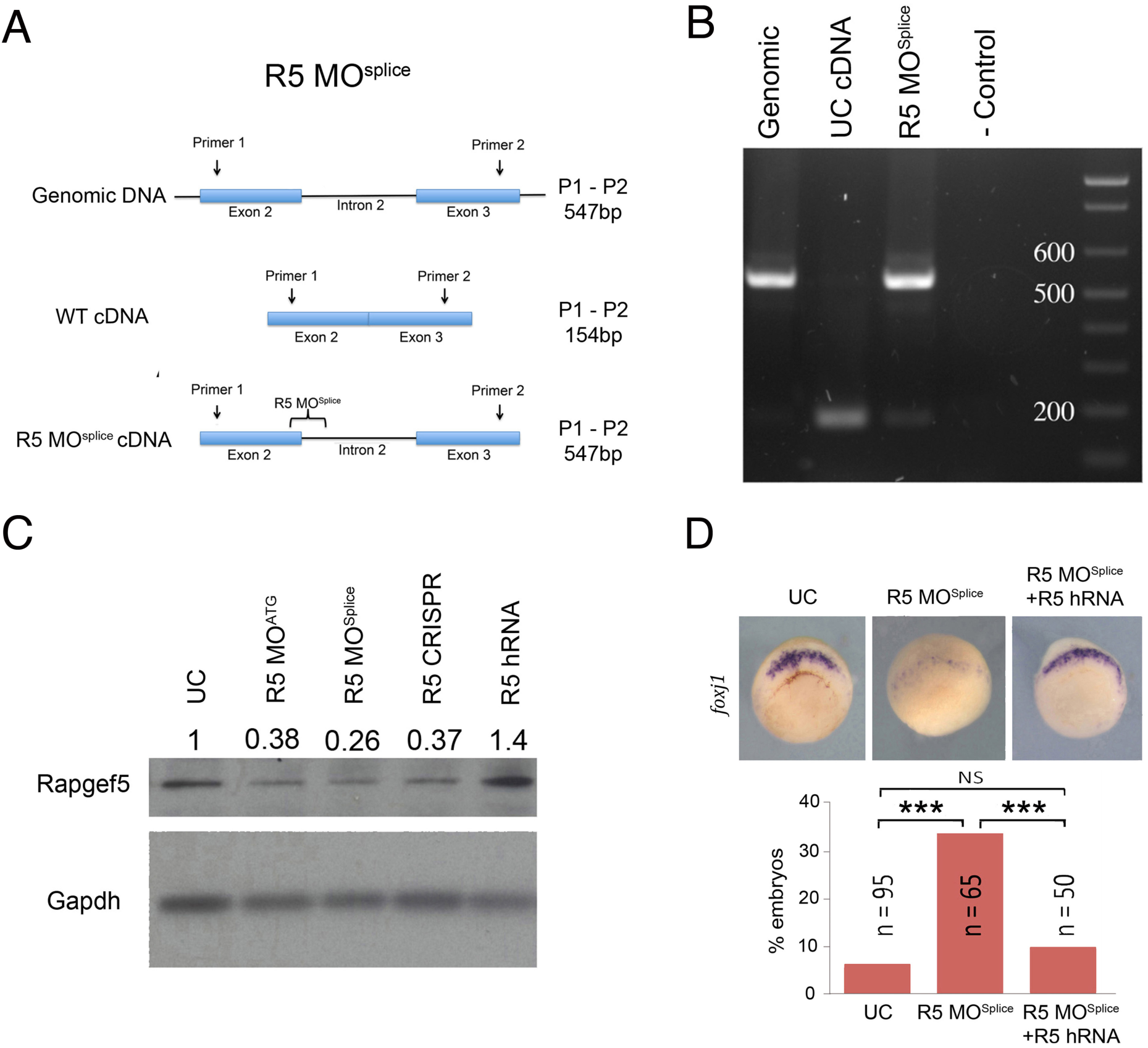
Related to Fig. 1; Depletion of Rapgef5. (A) Rationale of R5 MO^Splice^ morpholino design. The splice morpholino targets the splice donor site at the start of intron 2. Failure to excise intron 2 results in a premature stop codon early in the Rapgef5 protein. (B) RT-PCR confirmation that R5 MO^Splice^ causes retention of intron 2. In genomic DNA intron 2 spanning primers produce a band of 547bp. In WT cDNA lacking intron 2 they produce a band of 157bp. cDNA from R5 MO^Splice^ morphants has a strong band at 547bp indicating retention of intron 2 in mRNA. A no template negative control is also included. (C) Western blot demonstrating the efficacy of the R5 MO^ATG^ and R5 MO^Splice^ morpholino, and CRISPR mediated Rapgef5 depletion in stage 10 *Xenopus* embryos. Over expression of human RAPGEF5 mRNA increases protein levels. (D) Co-injection of human RAPFEG5 mRNA can rescue the loss of *foxj1* expression in Rapgef5 morphants, demonstrating the specificity of the morpholino. A triple asterisks indicate P<0.005.

**Figure S2.**
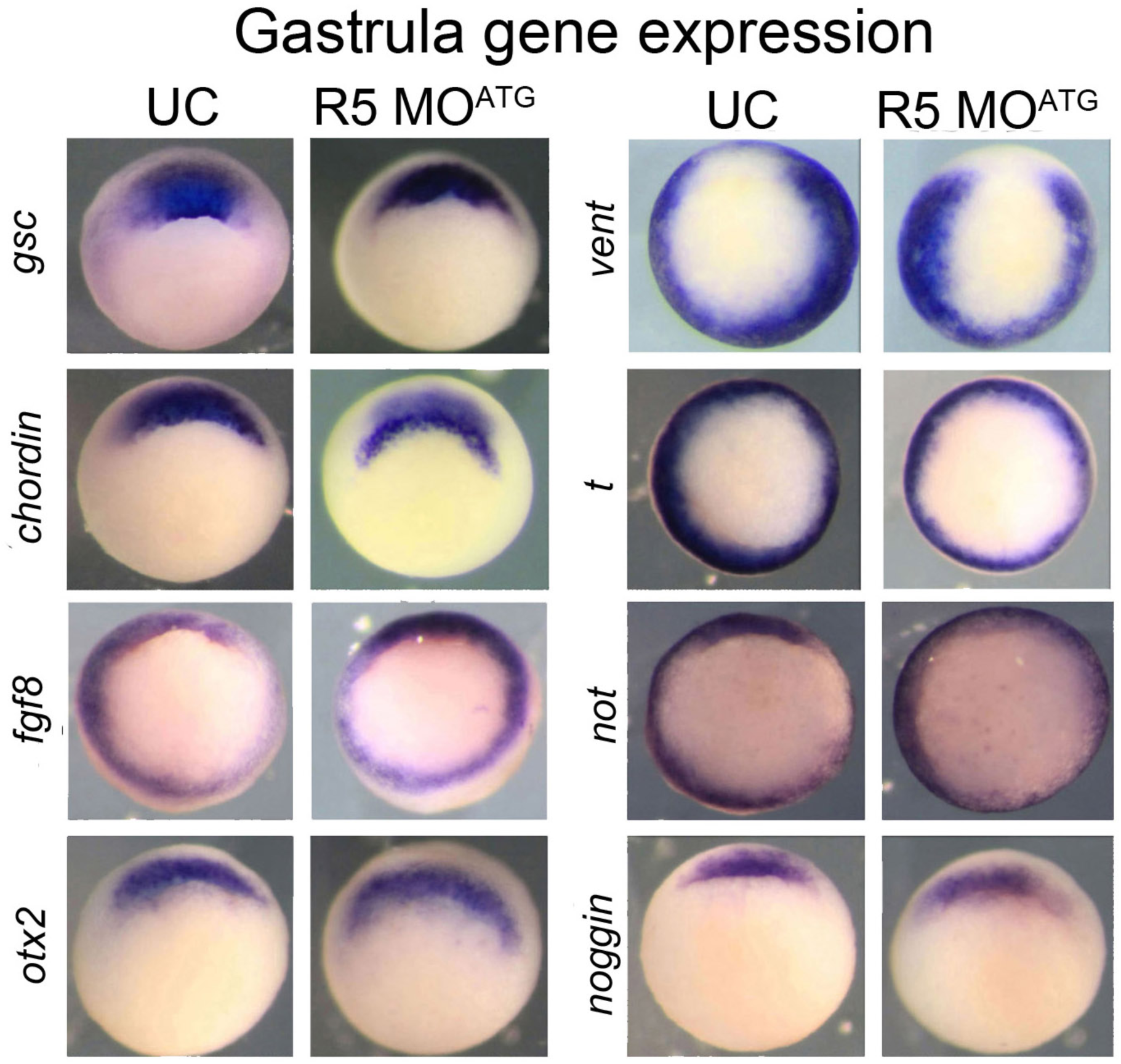
Related to Fig. 2; Expression of gastrulation genes in rapgef5 morphants. (A) Expression of numerous gastrulation markers (*gsc, chordin, fgf8, otx2, vent, t, not* and *noggin*) are not affected at stage 10 by depletion of Rapgef5. Vegetal views with dorsal to the top.

**Figure S3.**
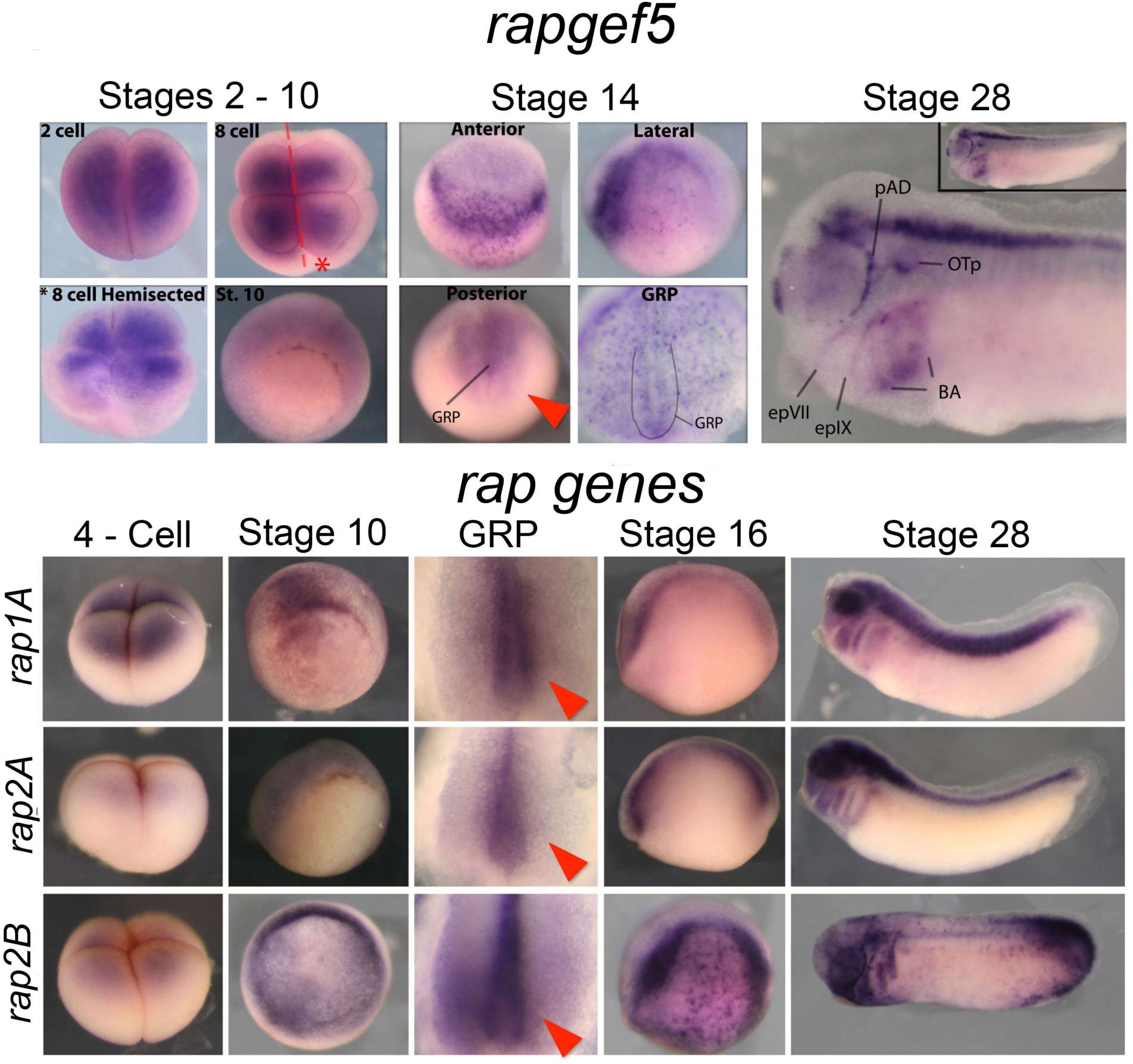
Related to Fig. 6; Developmental expression of Rapgef5 and Rap genes. *In situ* hybridization reveals *rapgef5* mRNA is present from the earliest stages of development. It is detected in the animal pole at the 2 cell and 8 cell stages. In the stage 10 embryos expression appears diffuse around the blastopore. At stage 14 transcripts are detected in the anterior neural folds and forming GRP. *rapgef5* expression is restricted to the anterior neural tube, brain, pharyngeal arches and cranial placodes at stage 28. Rap1A, Rap2A and Rap2B are all expressed in the 4 cell embryo, in the gastrulating stage 10 embryo and in the GRP at stage 16. Expression of all three genes is also strong in the neural folds at stage 16, while Rap2B also displays an interesting speckled pattern on the lateral ectoderm. All three are expressed in the neural tube, brain, eye and pharyngeal arches at stage 28.

**Figure S4.**
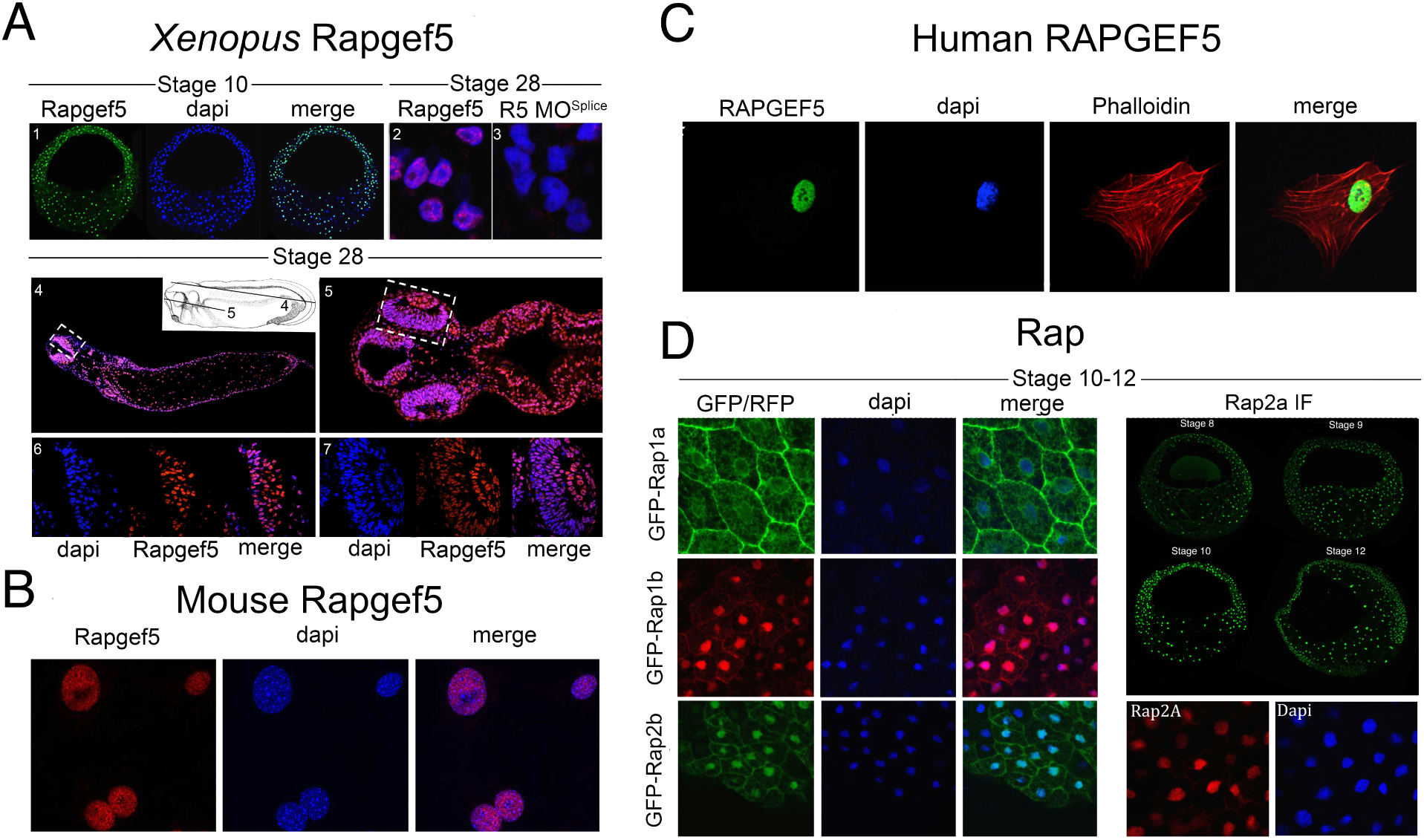
Related to Fig. 6; The subcellular localization of Rapgef5 and Rap proteins. (A1) Section of a stage 10 *Xenopus* embryo revealing that Rapgef5 is localized to the nuclei. (A2 - 7) Rapgef5 is detected specifically in the nuclei of sectioned paraffin embedded stage 28 *Xenopus* embryos but is reduced in morphants (Compare A2 and A3). The lines on the schematic in A4 represent the plane of section shown in A4 and A5. A6 and A7 are higher magnification of the white boxes in A4 and A5, respectively. (B, C) Rapgef5 localizes specifically to the nuclei of mouse MEFs and human RPE cells. (D) With the exception of Rap1A all rap proteins localize both to the plasma membrane and nucleus as assayed by injection of GFP/RFP tagged constructs or by immunofluorescence.

## STAR METHODS

### Contact for Reagent and Resource Sharing

Further information and requests for reagents may be directed to, and will be fulfilled by the corresponding author Mustafa K. Khokha

### Experimental Model and Subject Details

#### Xenopus

*X. tropicalis* were maintained and cared for in our aquatics facility, in accordance with Yale University Institutional Animal Care and Use Committee protocols. Embryos were produced by *in vitro* fertilization and raised to appropriate stages in 1/9MR + gentamycin as per standard protocols (del Viso and Khokha, 2012).

#### Mice

Production of the mouse *Catnblox(ex3)* allele, carrying a loxP flanked β-catenin exon 3, was as previously described(Harada et al., 1999). Heterozygous *Catnblox(ex3)* female mice were mated to a pCAGGCre-ER^TM^ male (Hayashi and McMahon, 2002) to produce tamoxifen inducible *pCAGG^+/cre^;Catnb^+/lox(ex3)^* embryos. Murine embryonic fibroblasts (MEFs) were obtained by standard protocols. All mice were of mixed genetic background and housed in the New Hunt’s House Biological Services Unit at King’s College London and all work was approved by the Ethical Review Board at King’s College London and performed in accordance with United Kingdom Home Office License 70/7441.

### Method Details

#### Morpholino oligonucleotides, mRNA and CRISPRs

Injections of *Xenopus* embryos were carried out at the one-cell stage as previously described (Khokha et al., 2002). We obtained ATG blocking (R5 MO^ATG^, 5’ – CTTTAAAAGTCTGACACCCATGAGC– 3′) or splice blocking (R5 MO^Splice^, 5’ – GCCTAAGGAAAAACTCTTACCTCCA – 3′) morpholino oligonucleotides from Gene Tools LLC and injected 8ng at the one cell stage to deplete Rapgef5. CRISPR sgRNAs containing the following rapgef5 target sequences (5’ – GGACTTCCTTCTCACATACA - 3’ and 5’ – GGGGACCCCGGAAAAGATTC - 3′) were designed from the v7.1 model of the *Xenopus* genome. sgRNAs and Cas9 protein was used to genetically knockdown *rapgef5* in F0 embryos as previously described (Bhattacharya et al., 2015). Full length human RAPGEF5 (clone HsCD00511998) was subcloned into the pCSDest2 vector using Gateway recombination techniques. Capped mRNAs were generated *in vitro* using the mMessage machine kit (Ambion) following the manufacturer’s instructions. Β-catenin-GFP (Addgene #16839), Stabilized β-catenin-GFP (Addgene #29684), NLS β-catenin-GFP (Addgene #29684), NLS-mCherry (Addgene #49313), TOPFlash (#12456) and FOPFlash (#12457) plasmids were obtained from Addgene. GFP-Rap1A, GFP-Rap1B, mCherry-Rap1B, GFP-XRap2, GFP-Rap2B and GFP-^RBD^RalGDS plasmids were generous gifts from Prof. Philip Stork at OHSU, Prof. Han at the Pohang University of Science and Technology, South Korea and Prof. Mark Philips at NYU.

#### Cardiac Looping

Stage 45 *Xenopus* embryos were paralyzed with benzocaine and scored with a light microscope. Looping was determined by position of the outflow tract. A D-loop was defined as the outflow tract going to the right, an L-loop was to the left, and an A-loop was midline.

#### RT-PCR

The R5 MO^Splice^ morpholino was targeted to the splice donor site in intron 2 of *rapgef5*. Retention of intron 2 was confirmed by RT-PCR, using the following primers: Forward:5’ TCCACTGTACAAGTGAAGGAAGA 3′, reverse: 5’ TGTCTTGTACCTCATCCAGG 3′.

#### *In Situ* Hybridization

Digoxigenin-labeled antisense probes for *chordin*: Tneu011C22, *coco:* TEgg007d24, *fgf8: Tneu007i10, foxj1:* Tneu058M03, *gdf3:* Tgas137g21, gsc: TNeu077f20, *noggin*: TNeu122a14, *not*; TEgg044k18, *otx2*; TGas114a06, *pitx2*; TNeu083k20, *Rap1A*: TEgg114f09, *Rap2A*: TGas143d22, *Rap2B*; TGas123a15, *t*; TGas116l23, *rapgef5;* IMAGE:6494175, *vent*; TNeu119b08, *xnr1* TGas124h10, and *xnr3;* Tgas011k18 were *in vitro* transcribed with T7 High Yield RNA Synthesis Kit (E2040S) from New England Biolabs. Embryos were collected at the desired stages, fixed in MEMFA for 1–2 hours at room temperature and dehydrated into 100% ETOH. Whole mount *in situ* hybridization was performed as described previously (Khokha et al., 2002).

#### Immunofluorescence

*Xenopus* embryos and mammalian cells were collected at desired time points and fixed in 4% paraformaldehyde/PBS (*Xenopus* = overnight (ON) at 4^o^C, cells = 15 mins at RT). Stage 10 *Xenopus* embryos were also embedded in paraffin and cut into 10 μm sections. Paraffin was removed using histoclear before beginning the IF. All samples were washed three times in PBS + 0.1% TritonX-100 before incubating in PBS + 0.1% Tween-20 + 3% BSA blocking solution for 1 hr at RT. Samples were then placed in blocking solution + primary antibody ON at 4^o^C. Samples were washed three times in PBS + 0.1% TritonX-100 before incubating in blocking solution + secondary antibody for 2hrs at RT. Samples were washed three times in PBS + 0.1% Tween-20, one of which included a 5 min incubation in dapi (1:5000), and washed again. Anti-RAPGEF5 (abcam, ab129008, 1:1000 dilution) was used as primary antibody. Alexa 488, 594, 647, and Texas red conjugated anti-mouse and rabbit secondary antibodies were obtained from Thermo Fisher Scientific. Alexa647 phalloidin (Molecular Probes, 1:40) was also used to stain cell membranes in some cases. Mammalian cells and *Xenopus* sections were mounted in Pro-Long Gold (Invitrogen) before imaging on a Zeiss 710 confocal microscope.

#### Mammalian cell culture and siRNA knockdown of Rapgef5

Mammalian embryonic fibroblasts (MEFs) were grown in DMEM supplemented with 10% FBS, L-glutamine, Penicillin streptomycin, hepes and β-mercaptoethanol. To knockdown Rapgef5, cells were transfected with non-overlapping Rapgef5 siRNAs (Santa Cruz # sc-152703), or with control siRNA (Santa Cruz sc-36869) using lipofectamine RNAiMAX transfection reagent (Thermo Fisher Scientific #13778100) as per the manufacturer’s instructions. Cells were incubated with siRNA overnight in serum free medium before being returned to MEF media for 36 hours.

#### TOPFlash Assays

*Xenopus* embryos were injected with 200pg of GFP-WT-β-catenin or GFP-Stabilized-β-catenin plasmids, along with 100pg of either the TOPFlash or FOPFlash (negative control) plasmids at the one cell stage. 20pg of Renilla plasmid was co-injected as an internal control. A subset of TOPFlash and β-catenin injected embryos were also injected with either 8ng of the Rapgef5 splice morpholino or 200pg of Gsk3 mRNA. Pools of 10 embryos were collected and lysed in 100μl passive lysis buffer.

In the mammalian cell culture experiments, Rapgef5 was depleted in *pCAGG^+/cre^;Catnb^+/lox(ex3)^* MEFs (carrying a tamoxifen inducible β-catenin gain of function allele) by siRNA knockdown. β-catenin was stabilized by the tamoxifen induced deletion of exon 3 after 36 hours and cells were collected for analysis at 48hrs. Luciferase was quantified using the Dual Luciferase Reporter Assay System (Promega #E1910) as per the manufacturer’s instructions. Each experiment was carried out in technical and biological triplicate.

#### Double axis assay and β-catenin localization experiments

For the secondary axis assay, embryos were injected at the one cell stage with 150pg of WT β-catenin-GFP, Stabilized β-catenin-GFP or NLS β-catenin-GFP. A subset of each group was then injected with either 8ng of the R5 MO^ATG^ morpholino or with full length GSK3 mRNA. Embryos were collected at stage 19 and scored for the presence of a secondary axis. To examine the subcellular localization of β-catenin in *Xenopus* embryos, 150pg of WT β-catenin-GFP, Stabilized β-catenin-GFP or NLS β-catenin-GFP were co-injected with NLS-cherry mRNA at the one cell stage. Samples were collected at stage 10 and fixed in 4% PFA overnight. The dorsal blastopore lip was dissected, mounted on coverslips in Pro-Long Gold (Invitrogen) and imaged with a Zeiss 710 confocal microscope. Image analysis was carried out using ImageJ.

#### Pharmacological inhibition of Gsk3

To inhibit GSK3α/β function during anterior development in WT *Xenopus* embryos and morphants, 6-bromoindirubin-3′-oxime (BIO, Sigma-Aldrich B1686) was added to the media at a concentration of 10 μM between stages 8 and 11. Media was then changed and the embryos allowed to develop. Anterior development was scored as normal, reduced or absent as presented in Fig. 2. For the nuclear/cytoplasmic fractionation experiments BIO was added at stage 4 (1 μM concentration) and the embryos were collected at stage 10.

#### Co-Immunoprecipitation and active Rap pulldown

Pools of stage 10 *Xenopus* embryos were lysed in NP-40 buffer (150 mM NaCl, 1.0% NP-40, 50 mM Tris, pH 8.0, 10 μl/embryo) on ice. Samples were centrifuged for 10 mins at 13,000 rpm and the supernatant removed to a clean tube. Supernatant was then pre-incubated with protein G beads (Bio-Rad, # 1614023) for 1 hour, centrifuged to collect supernatant, and incubated with primary antibody (Anti-β-catenin H-102, Santa Cruz, sc-7199) overnight. Protein G beads were then added to the sample solution and incubated for 3 hrs at 4^o^C. Lysate was centrifuged and the supernatant discarded. The beads were washed and centrifuged with 0.1% PBS-Tween six times. 2x SDS loading dye was added to the beads and heated to 95^o^C for five mins. The sample was centrifuged again and the lysate moved to a clean tube. For the active Rap1 and Rap2 pull down experiments, the active rap detection kit, (Cell signaling Technologies #8818) was used as per the manufacturer’s instructions. Samples were analyzed with Western blot.

#### Nuclear/cytoplasmic protein fractionation

Pools of 30 embryos were collected at stage 10 for each treatment including controls in eppendorf tubes, (a maximium of 35 and minimum of 20 embryos can be used with the solution volumes outlined in this protocol, but the protein concentrations will vary). All centrifugation steps were done at 4^o^C. 1 ml of Buffer E1 (see below for details for E1, E2, E3) was added to the embryos, and they were homogenized using a pipette. Lysates were centrifuged at 13,000 rpm at 4^o^C for 5 min. Approximately 600-650 μl of the supernatant was transferred to a new eppendorf tube (avoiding the lipid fraction) and placed on ice (cytoplasmic fraction). The remaining supernatant was discarded, and any lipids were removed from the tube walls using a Kimwipe, talking care not to disturb the pellet (nuclear fraction).

The nuclear and cytoplasmic fractions were then purified. For the cytoplasmic fraction, 11 μl of buffer CERII from the NE-PER Nuclear and Cytoplasmic Extraction Reagents Kit (Thermo-Scientific #78833) was added and tubes were vortexed for 15 sec, incubated on ice for 5 min and centrifuged for another 5 min. The supernatant (~ 500 μl) was transferred to a clean tube again avoiding lipid contamination. This cytoplasmic fraction was stored at -80^o^C until use.

The nuclear fraction pellet was resuspended in 1 ml of buffer E1 by pipetting (some material might remain insoluble), incubating on ice for 5 min and centrifuging for 5 min. The supernatant was discarded following centrifugation and any lipids were removed using Kimwipes without touching the pellet. This washing step was then repeated. After the second centrifugation the pellet was resuspended in 1 ml of buffer E2 and centrifuged for 5 min. The supernatant was discarded. This step was repeated once. Subsequently, the nuclear pellet was resuspended in 500 μl of buffer E3 and centrifuged for 5 min. The supernatant was discarded and the pellet was finally resuspended in 150 μl of Buffer E3. This is the nuclear fraction.

Protein concentrations of the nuclear and cytoplasmic fractions were determined using DC Protein Assay (Biorad # 5000112). Protein fractions were stored at -80^o^C. All buffers were supplemented with proteinase inhibitor (Complete, Roche # 11697498001) and PMSF (0.2 mM final concentration) prior to use. Buffer E1 was also supplemented with DTT (1 mM final concentration).

Buffer compositions: ***Buffer E1***: 50 mM HEPES-KOH, 140 mM mL NaCl, 1 mM EDTA (pH 8.0), 10% Glycerol (v/v), 0.5% Igepal CA-630 (v/v), 0.25% Triton X-100 (v/v). ***Buffer E2*:** 10 mM Tris-HCl (pH 8.0), 200 mM NaCl, 1 mM EDTA (pH 8.0), 0.5 mM EGTA (pH 8.0). ***Buffer E3*:** 10 mM Tris-HCl (pH 8.0), 200 mM NaCl, 1 mM EDTA (pH 8.0), 0.5 mM EGTA (pH 8.0), 1% (v/v) Sodium deoxycholate (from a 10% [w/v] solution), 5% (v/v) Sodium N-lauroylsarcosine (from a 10% [w/v] solution).

#### Western blotting

For total embryo western blots, pools of 10 control and treated embryos were collected at stage 9, 10 or 12 and placed in 100 ul of 1 x RIPA buffer. Embryos were then crushed using a pestle and spun down twice to separate lysate from lipids and debris. Embryos for nuclear and cytoplasmic analysis fractionations were isolated as described above. Western blotting was carried out following standard protocols and 45 μg of protein was loaded in each lane in 4-12% Tris-Bis gels. Anti-β-catenin (Santa Cruz sc-7199, 1:1000 in 5% milk), anti-active-β-catenin (Millipore 05-665), anti-GAPDH (Ambion, AM4300 1:5000 dilution), anti-β-actin (cytoplasmic marker, santa cruz Sc-4778), anti-Histone H3 (nuclear marker, Abcam, ab1791; 1:2000 dilution), anti-RAPGEF5 (abcam, ab129008, 1:200 dilution), anti-Rap1 (Enzo, ADI-KAP-GP125, 1:250) and anti-Rap2 (Abcam #ab166785, 1:500) primary antibodies were used. Anti-mouse or anti-rabbit HRP conjugated secondary antibodies were used (Jackson Immuno Research Laboratories, 715–035–150 or 211–032–171 1:15000 dilution). Bands were quantified using ImageJ and relativized to controls. Injections and protein fractionations were repeated 3 times. Western blots images show representative results and graphs show the averages of the different replicates.

### Quantification and Statistical Analysis

For *Xenopus* experiments we estimated 20–25 samples per experimental condition were necessary for statistical significance given the magnitude of the changes expected. Typically, many more samples were obtained for each experiment except as noted in the figures. Statistical significance is reported in the figures and legends. In all figures, statistical significance was defined as P<0.05. A single asterisk indicates P<0.05, while double and triple asterisks indicate P<0.01 and P<0.005, respectively. In Figure 1, the *in situ* results were analyzed by chi-squared test or Fishers exact test as appropriate for sample size. In Figure 2 the double axis assay data was examined using fishers exact test, the BIO treatment by Chi-squared and the TOPFlash by t-test. For Figure 3 and Extended Data 3, a t-test (two-tailed, type two) was used to determine significance of the ratiometric analysis, and the western blot data was analyzed by Anova Sidak. In Extended Data 2 fishers exact test was used to analyze the *in situ* data.

